# GE-BiCross: A Hierarchical Bidirectional Cross-Attention Framework for Genotype-by-Environment Prediction in Maize

**DOI:** 10.64898/2026.03.10.710816

**Authors:** Shuchang Zhou, Weipeng Fang, Runing Gao, Ting Zhao

**Affiliations:** Zhejiang Provincial Key Laboratory of Crop Genetic Resources, Institute of Crop Science, Plant Precision Breeding Academy, College of Agriculture and Biotechnology, Zhejiang University, Hangzhou, China

## Abstract

Genotype-by-environment interactions are central to crop adaptation and yield stability, yet they remain difficult to model for robust prediction across heterogeneous environments. Although enviromic profiling has improved the characterization of dynamic field conditions, most existing genomic prediction methods adopt a late-fusion strategy that encodes genomic and environmental information independently before global integration, thereby limiting their ability to resolve fine-scale, context-dependent G x E effects. Here, we developed GE-BiCross, a hierarchical bidirectional cross-attention framework for maize prediction. GE-BiCross incorporates a dual-path feature extraction module to disentangle independent and cooperative effects, a tokenized bidirectional cross-attention module to enable reciprocal genotype-environment interaction learning, and a mixture-of-experts module to adaptively capture heterogeneous response patterns across environments. Using a large-scale dataset of approximately 360,000 observations from 4,923 maize hybrids evaluated in 241 environments, GE-BiCross consistently outperformed conventional genomic prediction, machine learning, and deep learning baselines across six agronomic traits. The greatest improvements were observed for environmentally responsive and genetically complex traits. In particular, GE-BiCross achieved an R2 of 0.672 for grain yield and 0.880 for grain moisture, significantly surpassing all comparison models. Ablation analyses demonstrated that the three core modules make distinct and complementary contributions to predictive performance.These results show that deep, bidirectional integration of genomic and enviromic information can substantially improve modeling of complex G x E interactions, providing a powerful framework for interpretable genomic prediction and climate-smart crop breeding.

## Introduction

Global climate change and the increasing frequency of extreme weather events severely compromise crop yield stability[1]. Consequently, developing climate-resilient cultivars with phenotypic plasticity and adaptability to dynamic environments has become a central challenge in modern breeding[2]. Agronomic traits are jointly shaped by genotype (G), environment (E), and their complex interactions (G×E)[3-5]. Crucially, G×E interactions not only drive phenotypic variation but also limit both the transferability of genetic effects across environments and the accuracy of breeding predictions[6].

Driven by advances in enviromics, envirotyping has emerged as the third core typing technology, alongside genotyping and phenotyping[7-9]. It enables the fine-grained characterization of spatiotemporally continuous environmental factors throughout the crop growth cycle. While previous studies show that composite environmental variables (e.g., photothermal time) explain phenotypic variation better than single-factor variables,[9,10] crop responses to these factors exhibit strong developmental stage-specific heterogeneity. Different growth stages often correspond to distinct physiological mechanisms. Although integrating environmental data improves predictive performance[11], most existing genomic prediction models rely on a late-fusion paradigm. In this “separate encoding followed by fusion” framework, genomic and environmental features are learned independently and interact only globally at the final stage. This approach is inadequate for capturing locus- and stage-specific G×E response patterns, and it fails to reflect the biological mechanisms underlying environmental regulation of gene expression.

To address these limitations, this study proposes GE-BiCross, a hierarchical multimodal learning framework designed to deeply couple genomic and enviromic data during representation learning. At its core, GE-BiCross employs a cross-attention mechanism to establish a heterogeneous query–key–value mapping, enabling selective retrieval and deep interaction across modalities. While cross-attention has proven effective in modeling condition-dependent interactions in other biomedical applications, for example, EZSpecificity[12] and XATGRN[13], its application here allows the framework to explicitly disentangle and integrate the stable and condition-dependent effects of genetic and environmental factors. This effectively characterizes how genetic contributions are reconfigured in dynamic environments. Furthermore, a mixture-of-experts (MoE) strategy is introduced to adaptively model heterogeneous environmental response patterns via dynamic routing.

Evaluated on a large-scale dataset comprising 4,923 maize hybrids across 241 environments, GE-BiCross significantly outperformed existing models in predicting multiple key agronomic traits[14]. Notably, for grain moisture, a trait highly sensitive to environmental variation, the coefficient of determination (R^2^) improved by 16.6%. Ultimately, this study establishes a robust technical foundation for developing biologically interpretable, climate-smart breeding models.

## Material and methods

### Dataset Overview

The final dataset used for modeling comprised approximately 360,000 valid observations, covering 241 trial environments and 4,923 hybrids[14]. Missing value rates for all six traits were below 5% and were imputed using the mean of the corresponding trait. The dataset was randomly partitioned by environment into training, validation, and test sets with a ratio of 8:1:1, ensuring that data from the same environment did not appear in more than one subset.

### Dual-Path Feature Extraction and Effect Decoupling

To finely extract intrinsic patterns from each modality, a parallel attention module with an identical structure is applied to the genotypic and environmental data. For an input feature **X**, this module extracts two complementary effect representations. Independent effects are captured by a multi-layer perceptron (MLP) that models point-wise contributions from individual features:

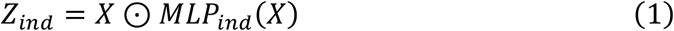

Concurrently, cooperative effects are captured by a multi-head self-attention (MHSA) mechanism that models global dependencies across feature dimensions:

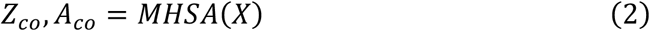

These two effect representations are then adaptively fused using a dynamic gating mechanism. The gate weights are generated dynamically from the cooperative features:

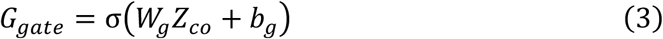

The final fused feature representation is:

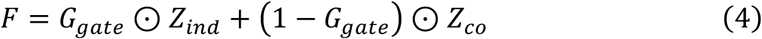

This process yields refined genotypic (*F*_*s*_)and environmental (*F*_*e*_) feature vectors.

### Tokenized Bidirectional Cross-Attention

To enable fine-grained interaction between genotype and environment, this module transforms the feature vectors into sequences of semantic units. Through learnable linear projections and reshaping operations, the genotypic and environmental features are mapped to token sequences 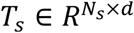 and 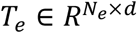, respectively. Here, *N*_*s*_ and *N*_*e*_denote the number of tokens for genotype and environment (set to 32 and 16 in this study), and *d* is the embedding dimension (set to 64). This tokenization allows the model to automatically cluster functionally related SNP groups or environmental factors into the same token.

Based on these sequences, bidirectional cross-attention is computed. Genotype-to-environment attention uses $T_s$ as queries and $T_e$ as keys and values:

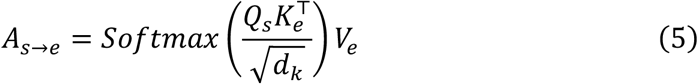

where *Q*_*s*_ = *T*_*s*_*W*_*Q*_, *K*_*e*_ = *T*_*e*_*W*_*K*_, *V*_*e*_ = *T*_*e*_*W*_*V* ○_ This mechanism aims to identify key environmental factors influencing a given genotype. Conversely, environment-to-genotype attention uses *T*_*e*_ as queries and *T*_*s*_ as keys and values:

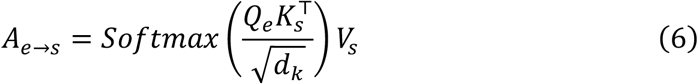

where 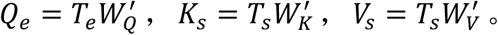 This aims to pinpoint genetic loci significantly activated under a specific environment. The outputs from each attention direction are average-pooled to yield *H*_*s*_ = *MeanPool*(*A*_*s*→*e*_) and *H*_*e*_ = *MeanPool*(*A*_*e*→*s*_), which are then concatenated and passed through a fusion MLP to produce the fused feature vector:

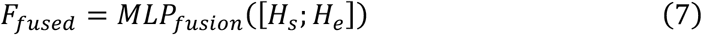

### Mixture-of-Experts Adaptive Prediction

To account for the heterogeneous response patterns inherent in G×E interactions (e.g., linear-, nonlinear-, or environment-specific dominance), an MoE layer is employed at the prediction stage. This layer consists of *k* parallel expert networks, each specializing in a distinct G×E pattern. A gating network dynamically computes expert weights based on the fused features and activates only the most relevant experts via a *Top* − *K* selection mechanism (set to *k* = 8 experts, *Top* − *K* = 2). The final output is the weighted sum of the selected experts’ outputs:

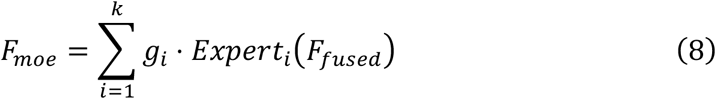

To prevent imbalanced expert utilization or collapse during training, an auxiliary load-balancing loss, *L*_*aux*_, is incorporated into the total loss function. This loss is based on the coefficient of variation of expert importance and load, following the approach detailed in Shazeer et al. (2017). (N. Shazeer, A. Mirhoseini, K. Maziarz, A. Davis, Q. Le, G. Hinton, and J. Dean, “Outrageously Large Neural Networks: The Sparsely-Gated Mixture-of-Experts Layer,” in *International Conference on Learning Representations (ICLR)*, 2017.)

### Experimental Setup

#### Data Splitting and Training Configuration

The dataset was randomly partitioned by environment into training, validation, and test sets with a ratio of 8:1:1. This splitting strategy ensures that data from the same environment do not appear in more than one subset, providing a rigorous evaluation of the model’s generalization performance to unseen environments. The model was trained using the AdamW optimizer with an initial learning rate of 1 × 10^−4^ and a cosine annealing scheduler. The batch size was set to 256. An early stopping mechanism was employed, halting training if the validation loss did not decrease for 20 consecutive epochs, with a maximum of 300 epochs. The total loss function is defined as:

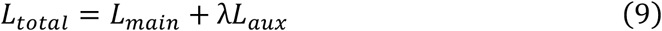

where the primary loss *L*_*main*_ is the Huber loss for robustness to outliers, and the auxiliary loss weight λ is set to 0.1.

#### Comparison Methods and Evaluation Metrics

The effectiveness of GE-BiCross was benchmarked against several genomic prediction methods, including Genomic Best Linear Unbiased Prediction (GBLUP), and traditional machine learning models like K-Nearest Neighbors (KNN), Random Forest (RF), and Gradient Boosted Trees (GBT). Additionally, we compared our model against GEFormer, a recent deep learning framework for G×E prediction.

Model performance was evaluated using two widely adopted metrics: the coefficient of determination (*R*^2), indicating the proportion of phenotypic variance explained by the model, and the Pearson correlation coefficient (PCC), measuring the linear correlation between predicted and observed values. All experiments were conducted on an NVIDIA GeForce RTX 5070ti GPU, and results are reported as the average of five independent runs to minimize stochastic variability.

## Result and Discussion

### Architecture of the GE-BiCross framework

We developed GE-BiCross, a hierarchical multimodal learning framework designed to explicitly model genotype-by-environment (G×E) interactions. As illustrated in Figure 1, the framework comprises three major components. First, a dual-path feature extraction and effect-disentanglement module captures genotype- and environment-derived representations while decomposing their main and interactive effects. Second, a tokenized bidirectional cross-attention module facilitates reciprocal information exchange between genetic and environmental features, enabling the fine-grained modeling of cross-modal dependencies. Third, a mixture-of-experts (MoE) adaptive prediction module dynamically integrates these learned representations to generate trait-specific predictions. The entire framework is trained end-to-end, allowing for the coordinated optimization of feature extraction, interaction modeling, and final prediction.

**Figure 1.**
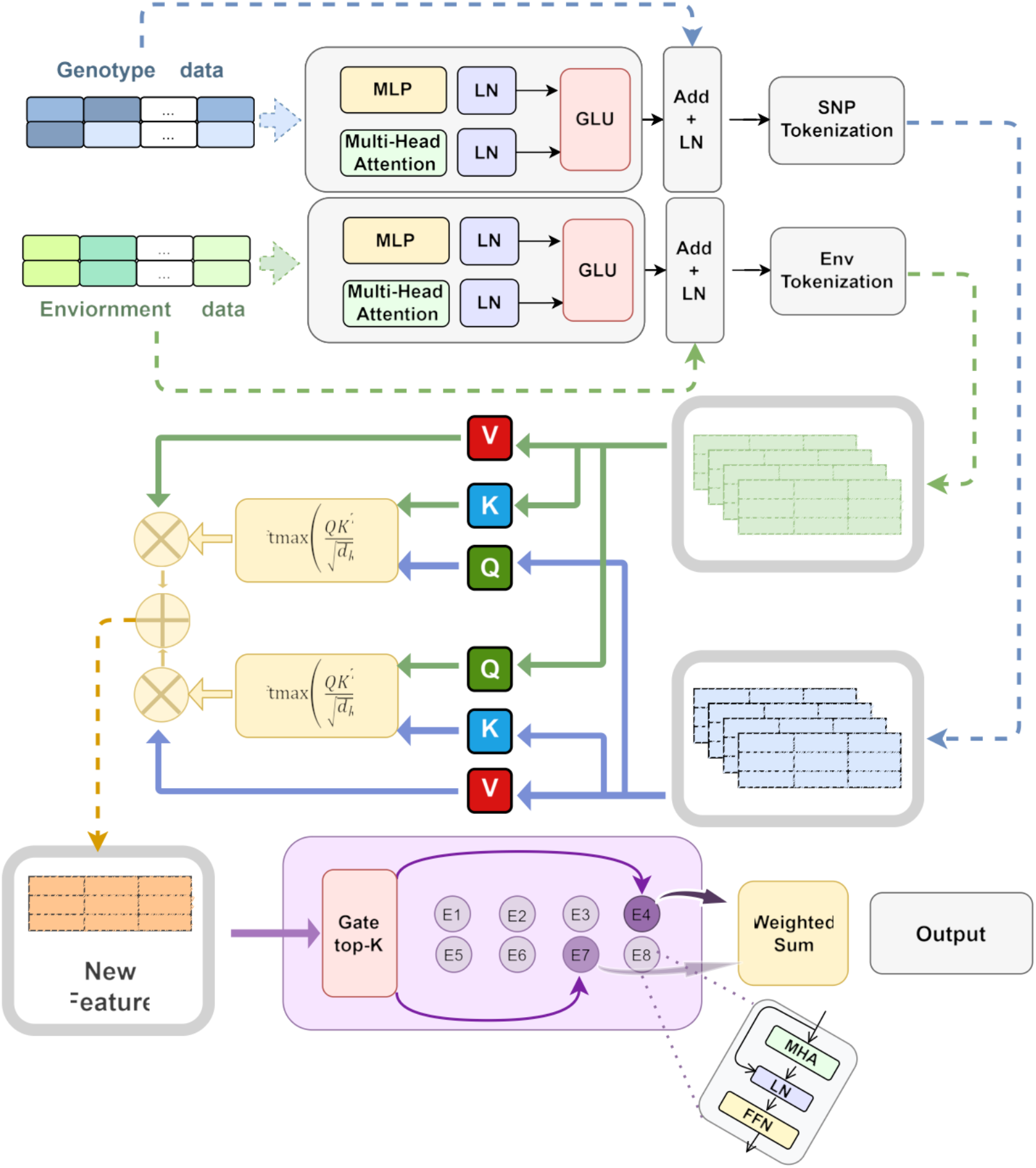
Schematic overview of the GE-BiCross architecture with a hierarchical mixture-of-experts (MoE) framework for genomic–environmental interaction modeling. The GE-BiCross model comprises three major components: (i) a genomic tokenization layer that transforms single nucleotide polymorphism (SNP) markers into position-aware embeddings, together with an environmental encoder for multi-modal environmental inputs, including meteorological, soil, and management variables; (ii) a bidirectional cross-attention module for the fusion of genomic and environmental representations; and (iii) a hierarchical MoE module with expert subnetworks and gating networks for conditional feature routing.

### GE-BiCross improves prediction across diverse agronomic traits

To evaluate its predictive performance, we compared GE-BiCross with representative conventional genomic prediction, classical machine learning, and deep learning methods (GBLUP, KNN, RF, GBT, and GEFormer) across six agronomic traits. GE-BiCross consistently achieved the highest predictive accuracy on the validation set for all six traits, demonstrating its superiority across traits with distinct genetic architectures and varying levels of environmental sensitivity.

The most substantial improvement was observed for yield, a highly complex trait strongly influenced by G×E interactions. GE-BiCross achieved an R^2^ of 0.672, outperforming the second-best model, RF (R^2^ = 0.615), by 0.057 (a 9.3% relative improvement) and surpassing GEFormer (R^2^ = 0.515) by 0.157 (a 30.5% relative improvement). Pairwise t-tests confirmed that these performance gains over all baseline models were statistically significant (p < 0.01). This highlights the capability of GE-BiCross to capture the complex interaction effects essential for accurate yield prediction.

A similarly strong advantage was observed for grain moisture, a trait highly responsive to environmental variation. GE-BiCross achieved an R^2^ of 0.880, compared with 0.849 for RF and 0.755 for GEFormer, yielding absolute improvements of 0.031 and 0.125, respectively. The improvement over RF was statistically significant (p < 0.05), while the improvement over GEFormer was highly significant (p < 0.001) (Figure 2). These results underscore the importance of explicit G×E modeling for environmentally sensitive traits.

**Figure 2.**
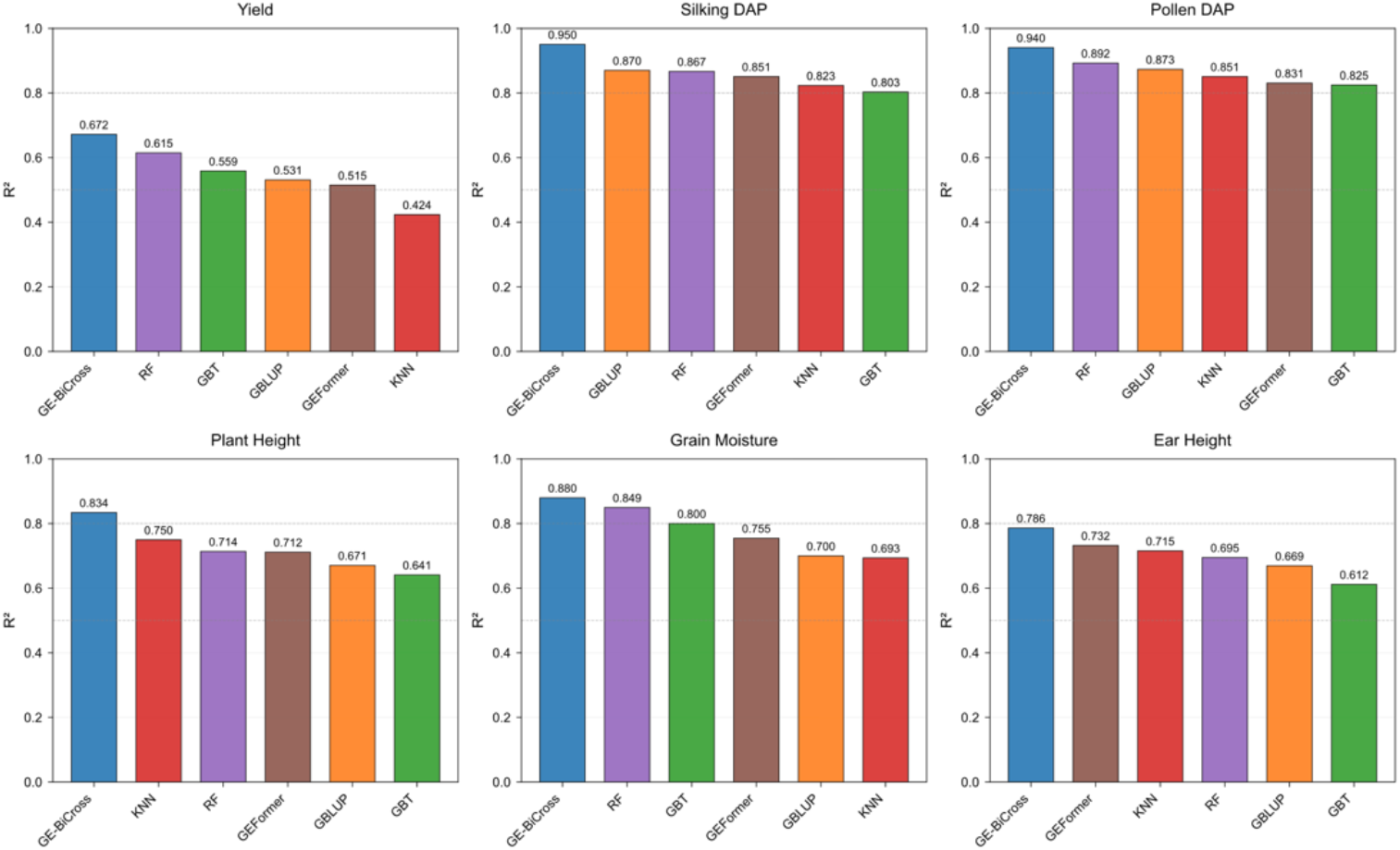
Performance comparison of genomic prediction models across six maize traits. The figure compares the predictive performance of six models, including GE-BiCross, Random Forest (RF), Gradient Boosting Trees (GBT), Genomic Best Linear Unbiased Prediction (GBLUP), GEFormer, and K-Nearest Neighbors (KNN), across six maize traits: grain yield, days to silking, days to pollen shed, plant height, grain moisture, and ear height. Predictive performance is evaluated using the coefficient of determination (R^2^).

**Figure 3.**
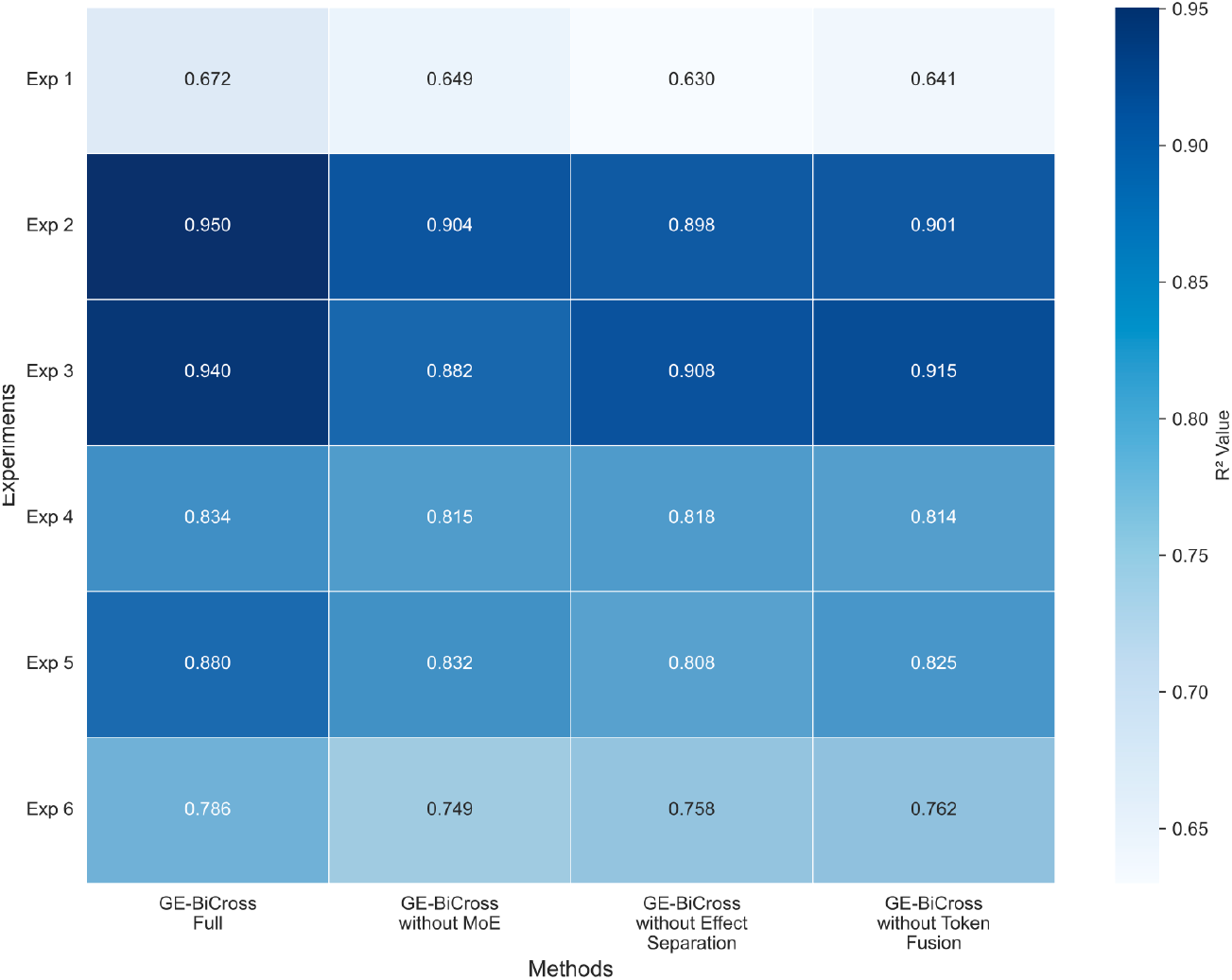
Ablation analysis of the GE-BiCross model across six maize traits. The heatmap shows the predictive performance of the full GE-BiCross model and three ablated variants: GE-BiCross without the mixture-of-experts (MoE) module, GE-BiCross without the effect separation module, and GE-BiCross without token fusion. Model performance is compared across six maize traits, including grain yield, days to silking, days to pollen shed, plant height, grain moisture, and ear height, to illustrate the contribution of individual architectural components.

GE-BiCross also delivered superior performance for flowering-related phenological traits. For silking date, the model achieved an R^2^ of 0.950, outperforming GBLUP (R^2^ = 0.870) by 0.080 (a 9.2% relative improvement). For pollen date, GE-BiCross reached an R^2^ of 0.940, exceeding RF (R^2^ = 0.892) by 0.048 (a 5.4% relative improvement). Although the baseline methods already performed well for these traits, the gains achieved by GE-BiCross remained statistically significant (p < 0.01). This indicates that the explicit integration of genetic and environmental information provides additional predictive value, even for traits with strong genetic determinism. Finally, GE-BiCross demonstrated consistent improvements for morphological traits.

The model achieved an R^2^ of 0.834 for plant height, outperforming KNN (R^2^ = 0.750) by 0.084 (an 11.2% relative improvement). For ear height, GE-BiCross attained an R^2^ of 0.786, exceeding GEFormer (R^2^ = 0.732) by 0.054 (a 7.4% relative improvement). Both performance gains were statistically significant (p < 0.05).

### Ablation analysis reveals distinct contributions of key GE-BiCross modules

To determine the contribution of each architectural component to predictive performance, we performed ablation analyses by selectively removing or replacing core modules of GE-BiCross. Specifically, we evaluated three reduced variants: GE-BiCross w/o MoE, in which the mixture-of-experts layer was replaced by a single fully connected layer; GE-BiCross w/o effect disentanglement, in which the dynamic gating mechanism separating independent and cooperative effects was removed and only cooperative features were retained; and GE-BiCross w/o token fusion, in which the bidirectional cross-attention module was replaced by direct concatenation of genotypic and environmental representations.

For grain moisture, the full GE-BiCross model achieved an R^2^ of 0.880, whereas all ablated variants showed substantial reductions in predictive accuracy. The largest decline was observed when the MoE module was removed (R^2^ = 0.808), corresponding to an absolute reduction of 0.072 (8.2% relative decrease). Replacing token fusion with simple concatenation reduced performance to R^2^ = 0.825 (absolute decrease of 0.055, 6.3% relative), whereas removing effect disentanglement resulted in an R^2^ of 0.832 (absolute decrease of 0.048, 5.5% relative). All differences between the full model and the ablated variants were highly significant (p < 0.001), indicating that each module contributes substantially to prediction of this environmentally sensitive trait, with the MoE module having the strongest effect.

A different pattern was observed for yield, for which the full model achieved an R^2^ of 0.672. In this case, the most pronounced performance loss resulted from removal of the effect-disentanglement module, which reduced the R^2^ to 0.630 (absolute decrease of 0.042, 6.3% relative). Removing token fusion reduced performance to R^2^ = 0.641 (absolute decrease of 0.031, 4.6% relative), whereas removal of the MoE module had a comparatively smaller effect (R^2^ = 0.649; absolute decrease of 0.023, 3.4% relative).

The reduction caused by eliminating effect disentanglement was statistically significant (p < 0.01), suggesting that explicit separation of independent and cooperative effects is particularly important for prediction of yield, a trait characterized by complex G×E architecture.

For flowering-related traits, the contributions of individual modules were more evenly distributed. In prediction of silking date, the full model reached an R^2^ of 0.950, whereas the variants without MoE, effect disentanglement, or token fusion achieved R^2^ values of 0.904, 0.898, and 0.901, respectively. These correspond to relative decreases of 4.8%, 5.5%, and 5.2%, indicating that all three components contribute comparably to predictive performance. The differences between the full model and all ablated variants were highly significant (p < 0.001), supporting the conclusion that robust prediction of flowering phenotypes requires coordinated contributions from adaptive expert routing, effect disentanglement, and cross-modal feature integration.

Taken together, these ablation results demonstrate that the three major modules of GE-BiCross make nonredundant and trait-dependent contributions to model performance. The MoE module appears to be particularly important for environmentally responsive traits such as grain moisture, whereas effect disentanglement is especially critical for complex traits such as yield. By contrast, for flowering traits, predictive gains arise from the complementary action of all modules. These results provide mechanistic support for the overall design of GE-BiCross and indicate that its performance advantage is not attributable to a single architectural component.

## Acknowledgements

This study was supported by the Biological Breeding-National Science and Technology Major Project (2023ZD04076).

